# Revisiting Mitchell’s chemiosmotic theory in light of new stoichiometric reaction equations of ATPase II

**DOI:** 10.1101/615880

**Authors:** Arindam Kushagra

**Author notes:** Corresponding Author: Arindam Kushagra.

## Abstract

Chemiosmotic theory has been reining the field of bioenergetics since its inception. As stoichiometric mechanisms of the underlying chemical reactions have now been elucidated, it calls for a revision in the relationships derived by Mitchell in his seminal review paper. Here in this work, new formulation of the relationship between the *pH* difference and transmembrane potential, as derived in light of modified ATPase II reaction stoichiometry, has been proposed for the first time. This formulation results in accurate estimation of dependence of transmembrane potential on the *pH* difference across the two sides of the mitochondrial membrane. Thus, this work is of potential interest and will enable researchers working in the field of bioenergetics involving chemiosmotic theory to come-up with more exact mechanistic explanations.

## Introduction

ATP is known to be the energy currency of the living cell. Dephosphorylation reactions within the cells provide chemical energy required for sustenance of biochemical processes essential to living organisms, with ATP breaking into ADP and inorganic phosphate, P_i_. Cellular ATP is synthesised with the help of F-type ATP synthase in many organisms, which works on a proton motive force (p.m.f.) that derives energy from the concentration gradients of H^+^ ions across the membranes along which ATP metabolism takes place. Usually, such site resides inside the mitochondria.

A notable work has been done in this regard by Peter Mitchell proposing vectorial H^+^ ion transport across the mitochondrial membranes. His work discusses generating an equivalent amount of p.m.f. which is further utilized for the metabolism (synthesis/hydrolysis) of ATP with the help of transmembrane enzymes like F-type ATP synthase/ATPase. In the 1966 review paper [1], a simplistic empirical equation for ATP synthesis was used with coefficients all the reactants as well as products were unity, which is usually not the case for any multi-molecular complex chemical reactions. Current work aims to modify Mitchell’s formulation of equations (7) to (14) in his review paper in light of new reaction mechanisms that have now been established, as discussed further. **Here, the author wishes to point out that the definition of *pH* used in this paper is the same that was prevalent when Mitchell’s review paper was first published in 1966 i.e. *pH* = −log_10_[H^+^]. The currently accepted definition, *pH* = −log_10_{H^+^}, was approved as a norm by IUPAC in 1979 [2]**. It may be noted that the electrochemical activity {} is given by the product of *γ*and [], where *γ* represents the activity coefficient of the respective ionic species and [] represents the ionic concentration i.e. {} = *γ*[]. The formulation presented in this paper for the first time lead to the estimation of electrochemical potential in very close approximation of the Nernst resting membrane potential at zero external transmembrane voltage.

## Formulation of modified relationship

The accepted mechanism of ATP hydrolysis/synthesis pertinent to chemiosmotic theory considers the hydrolysis of ATP to form ADP and inorganic phosphate species, which is assumed to be just reverse of the ATP synthesis from the same species. The revised stoichiometric equation of ATP synthesis from ADP [3] which forms the basis of this work is given by reaction R1:

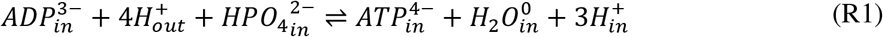

The above ATP synthesis equilibrium can be represented in terms of electrochemical activity, denoted by curly brackets {}, as shown in equation (1):

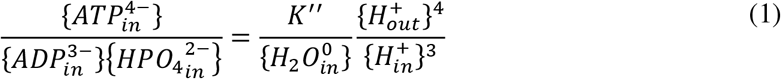

The values of activity coefficients, *γ*, may be calculated by Davies’ equation (used for ionic strength, I<0.5M) [4], as given by following relationship:

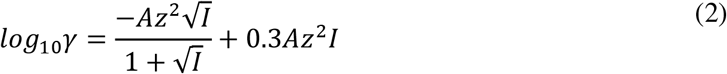

where, *A* = 0.5, *z* = charge on the ion & ionic strength 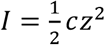 (*c* is concentration of the ion).

If there is no external potential applied across the membrane, the resulting electrochemical potential will be just due to the difference in *pH* values on either side of the membrane. These potential values are also known as resting membrane potential in the scientific parlance.

At this juncture, the author proposes a novel relationship between the activity coefficients (*γ*), electrochemical potentials of hydronium ions {H^+^} on either side of the membrane at the site of ATP synthesis, difference in *pH* values, transmembrane potentials due to different pH values (given by *ΔE_pH_*) & due to presence of other ions like K^+^ (given by *ΔE_ion_*) and electrochemical potentials due to those ions (given by *EP*), as shown in equation (3):

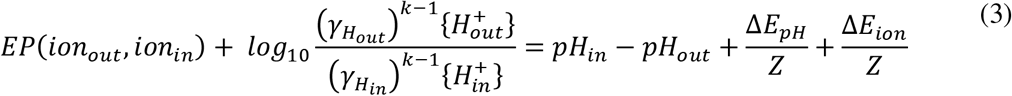

In the equation (3), *ΔE* represents the transmembrane potential (in mV) and 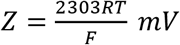 (*R*: Universal gas constant, *F*: Faraday’s constant, T: Temperature in K).

When the review was first published in 1966, the accepted definition of *pH* was

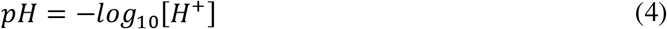

It was later modified in 1979, to

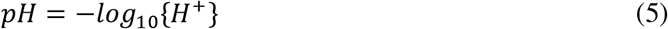

Here the author wishes to make a cautionary remark that since Mitchell’s work was published 13 years before the current definition of pH was accepted by IUPAC, it would be unjust to judge the accuracy (or the lack of it) of Mitchell’s formulation while viewing them from the perspective of equation (5). To calculate the values of activity coefficients from Davies’ equation (equation (2)), concentrations of the species involved in F-ATP synthase reaction (for ATP synthesis) have been given in the literature [3] as shown in Table 1:

**Table 1.** Experimentally determined concentrations of species involved in ATP synthesis reaction using F-ATP synthase (respective values have been taken from Supplementary Information section of [3])

| *Ionic species* | *Concentration (M)* |
| --- | --- |
| $ATP^{4-}$ | $2 \times 10^{-4}$ |
| $HATP^{3-}$ | $7.96 \times 10^{-4}$ |
| $H_2ATP^{2-}$ | $3.79 \times 10^{-6}$ |
| $ADP^{3-}$ | $1.99 \times 10^{-4}$ |
| $HADP^{2-}$ | $3 \times 10^{-4}$ |
| $H_2ADP^{-}$ | $6.85 \times 10^{-7}$ |
| $HPO_4^{2-}$ | $3.77 \times 10^{-3}$ |
| $H_2PO_4^{-}$ | $6.23 \times 10^{-3}$ |
| $H_2O^0$ | 1.00 |
| $H_{in}^{+}$ | $1 \times 10^{-7}$ |
| $H_{out}^{+}$ | $3.16 \times 10^{-7}$ |

From the values given in Table 1 and from Davies’ equation, we obtain the activity coefficients for the species participating in the ATP synthesis as shown in Table 2:

**Table 2.** Values of activity coefficients calculated using the concentrations given in Table 1 and Davies’ equation

|  |  |
| --- | --- |
| $-\log_{10}\gamma_{ATP}$ | 0.2949 |
| $-\log_{10}\gamma_{ADP}$ | 0.1270 |
| $-\log_{10}\gamma_{HPO_4}$ | 0.145 |
| $-\log_{10}\gamma_{H_{out}}$ | $1.99 \times 10^{-4}$ |
| $-\log_{10}\gamma_{H_{in}}$ | $1.118 \times 10^{-4}$ |

As per the definition of electrochemical activity, {} = *γ*[], equation (3) may be rewritten as,

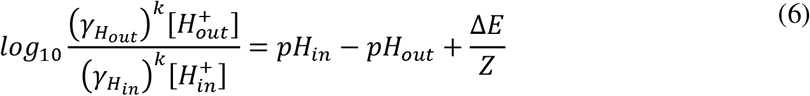

Modifying and rearranging equation (6) to derive a relationship between the activity coefficients and the transmembrane potential at steady state, we get the following relationship after reduction:

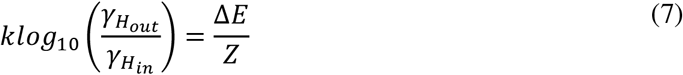

where, *k* represents the proportionality constant when only resting membrane potential resulting due to *pH* difference across the membrane, is considered. The calculated values of parameter *k*, in the above equation have been observed to vary with *pH_in_ - pH_out_* as shown in Table 3:

**Table 3.** Variation of exponent of activity coefficients, *k* with *pH_in_ - pH_out_*

| $pH_{in} - pH_{out}$ | $k$ |
| --- | --- |
| 0.5 | 5 |
| 1 | 4 |
| 1.5 | 3 |
| 2 | 2 |
| 2.5 | 1.3 |
| 3 | 0.9 |
| 3.5 | 0.57 |
| 4 | 0.36 |

The data depicted in Table 3, is shown graphically in Fig.1. It follows a quadratic polynomial relationship between the parameter *k* and *pH_in_ - pH_out_*.

**Fig. 1.**
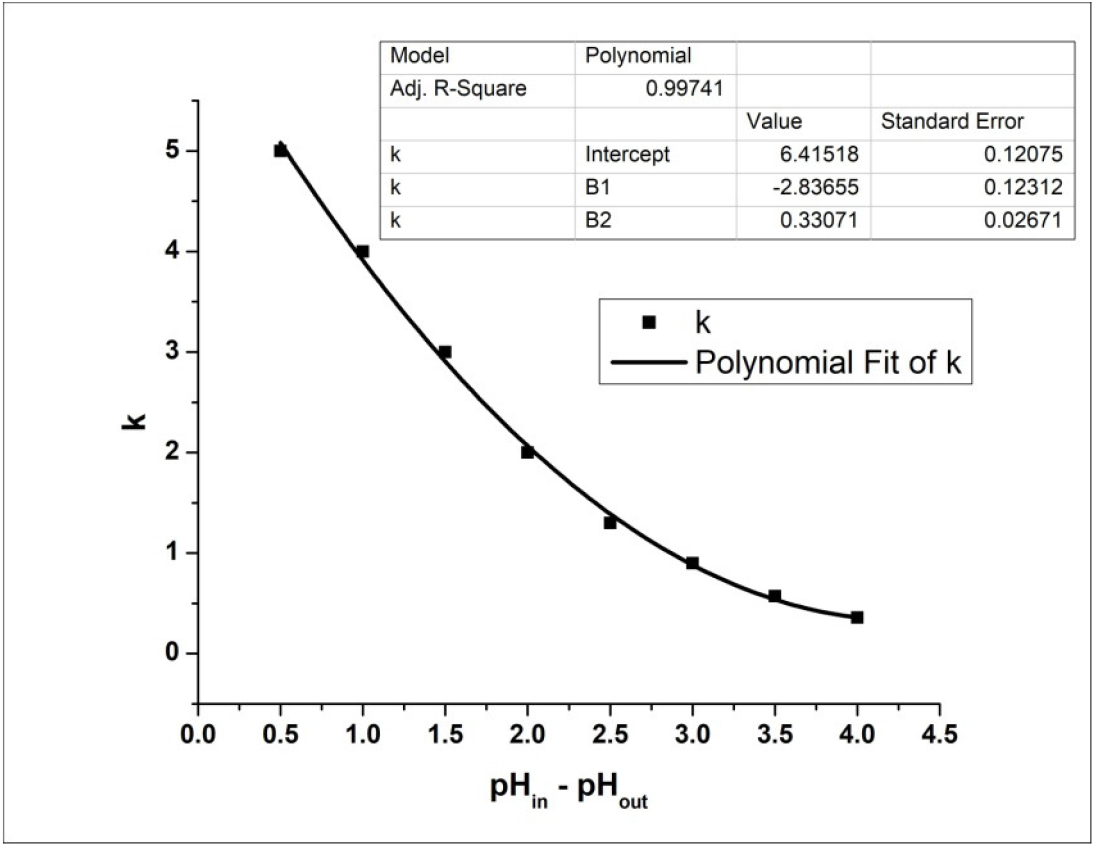
Relationship between parameter *k* and *pH_in_ - pH*_out_, showing a quadratic polynomial fit. The fitting parameters are shown in the “inset” table. The polynomial fit was done with Origin Pro8^®^.

Putting the “as calculated” activity coefficients from Table (2) and *Z* ≈ 60 *mV* in equation (7) and after putting *k* from Table 3, we get the values of *ΔE* which represents the resting membrane potential due to *ΔpH* (when no external transmembrane potential is applied).

To further investigate the applicability of the proposed relationship, as shown in equation (3) of this paper, we continue to derive new relationships between the transmembrane potential and *pH* difference. Consequently, taking log_10_ of equation (1) on both sides, we get the following relationship:

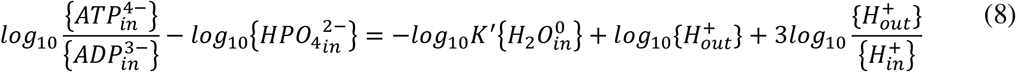

After putting the concentration values of respective species (from Table 1) and calculated values of log10*γ*in equation (8), we get the following relationship:

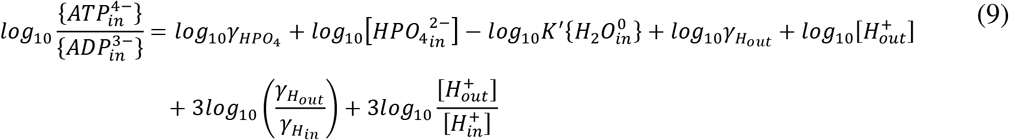

Substituting the value of 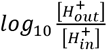 from equation (6), we arrive at the following relationship:

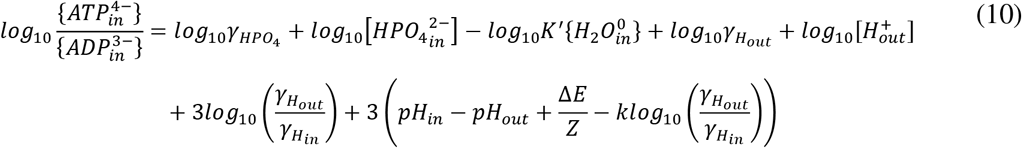

It turns out that the terms containing the activity coefficients of ionic species are very small in magnitude in comparison to the other terms in equation (9). The value of 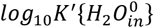 which is the hydrolysis constant for ATP in the above relationship can be approximated to 5 at 300 K [1]. Therefore, the final relationship is given by equation (11):

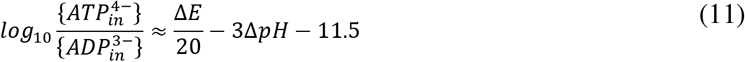

Here, the difference in *pH* values is defined as *ΔpH* = *pH_out_* - *pH_in_*. After suitable rearrangements, the above equation yields equation (12) as under:

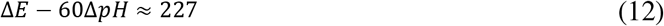

As per the experimental concentrations of participating ionic species during ATP synthesis, listed in Table 1, it is observed that the transmembrane potential from equation (12) comes out to be ≈197 mV. This is in accordance with the reported transmembrane potential from isolated mitochondria to be in the range of 180-220 mV, as discussed by Ramzan *et al*. [5]. Thus, we observe the accuracy of equation (12) in successfully predicting the transmembrane potential as a function of *pH* difference within physiological limits by using the proposed relationship as discussed in equation (3).

## Discussion

In the present work, a new formulation has been provided for chemiosmotic coupling in ATP metabolism to elucidate a modified dependence of transmembrane potential on the *pH* difference. These equations may further be used for varying *pH* conditions, inside and outside the membrane allowing H^+^ translocation for ATP metabolism, covering the entire biological range of *pH* difference across the membrane when ATP metabolism takes place. Specifically, the exponents of activity coefficients involved in the new proposed formulation have been shown to have a quadratic dependence (**R**^2^: 0.99741) on the *pH* difference across the membrane. Validity of the equations has been tested by calculating the mitochondrial transmembrane potential, which came out to be in perfect agreement with the values reported in the literature. This paper will be of immense interest to the scientific community working in the field of bioenergetics, especially oxidative phosphorylation and will help in developing a better understanding in the light of newly derived equations that are analogous to the ones reported by Mitchell in his review paper.

## Conflict of Interest

The author declares no conflict of interest.

